# Evidence for a Causal Dissociation of the McGurk Effect and Congruent Audiovisual Speech Perception via TMS

**DOI:** 10.1101/2023.11.27.568892

**Authors:** EunSeon Ahn, Areti Majumdar, Taraz Lee, David Brang

## Abstract

Congruent visual speech improves speech perception accuracy, particularly in noisy environments. Conversely, mismatched visual speech can alter what is heard, leading to an illusory percept known as the McGurk effect. This illusion has been widely used to study audiovisual speech integration, illustrating that auditory and visual cues are combined in the brain to generate a single coherent percept. While prior transcranial magnetic stimulation (TMS) and neuroimaging studies have identified the left posterior superior temporal sulcus (pSTS) as a causal region involved in the generation of the McGurk effect, it remains unclear whether this region is critical only for this illusion or also for the more general benefits of congruent visual speech (e.g., increased accuracy and faster reaction times). Indeed, recent correlative research suggests that the benefits of congruent visual speech and the McGurk effect reflect largely independent mechanisms. To better understand how these different features of audiovisual integration are causally generated by the left pSTS, we used single-pulse TMS to temporarily impair processing while subjects were presented with either incongruent (McGurk) or congruent audiovisual combinations. Consistent with past research, we observed that TMS to the left pSTS significantly reduced the strength of the McGurk effect. Importantly, however, left pSTS stimulation did not affect the positive benefits of congruent audiovisual speech (increased accuracy and faster reaction times), demonstrating a causal dissociation between the two processes. Our results are consistent with models proposing that the pSTS is but one of multiple critical areas supporting audiovisual speech interactions. Moreover, these data add to a growing body of evidence suggesting that the McGurk effect is an imperfect surrogate measure for more general and ecologically valid audiovisual speech behaviors.

## Introduction

While speech perception is predominantly an auditory process, face-to-face conversations benefit from concurrent visual cues that provide both complementary and redundant information about an auditory stimulus, particularly in noisy environments (Campbell, 2008; MacLeod & Summerfield, 1987; Sumby & Pollack, 1954). For example, listeners can derive both timing and phonemic information about spoken words simply by watching a speaker’s mouth movements (Luo et al., 2010; Plass et al., 2020; Schroeder et al., 2008). This visual augmentation can significantly improve speech perception accuracy, especially when acoustics are compromised. Given the inherent multisensory nature of speech, it is important to examine how the brain enables vision to support language to understand speech processing in naturalistic contexts.

Traditionally, studies of audiovisual processing have relied on the use of the McGurk effect, in which an auditory phoneme (e.g., /ba/) is paired with the visual movie from a different phoneme (e.g., /ga/), resulting in the perception of a fused or unique sound (e.g., /da/) (McGurk & MacDonald, 1976). Using McGurk stimuli, prior research has identified the left posterior superior temporal sulcus (pSTS) as a crucial region that facilitates the integration of auditory and visual information (Benoit et al., 2010; Bernstein et al., 2008; Irwin et al., 2011; Nath et al., 2011; Sekiyama et al., 2003; Szycik et al., 2012). For example, individual differences in the strength of the McGurk effect are correlated with fMRI activity in the pSTS during the experience of the illusion (Nath & Beauchamp, 2012), and both transcranial magnetic stimulation (TMS) and damage following a stroke in this region are associated with reduced McGurk effect percepts (Beauchamp et al., 2010; Hickok et al., 2018). However, researchers have recently questioned whether the findings from research using McGurk combinations can generalize to more natural audiovisual integration processes (for review see Alsius et al., 2018). For example, numerous studies (Brown & Braver, 2005; Hickok et al., 2018; Van Engen et al., 2017) have reported weak correlations between the McGurk effect and other measures of audiovisual speech processing in individuals, raising doubts about whether the mechanisms that enable the McGurk effect are the same as those that process natural audiovisual speech.

Whether the McGurk effect and more general audiovisual processes rely on the same mechanisms is a significant issue in the field because the McGurk effect had long been accepted as a standard measure for quantifying audiovisual speech integration (Alsius et al., 2018; Van Engen et al., 2022). This holds especially true for clinical populations including those with autism spectrum disorder (ASD). Individuals with ASD often exhibit difficulties in communication and social interaction and many researchers believe that this could be in part attributed to impairments in multisensory processing. To study this, researchers have repeatedly used the likelihood of McGurk percepts to demonstrate that individuals with ASD show altered multisensory processing (Feldman et al., 2022; Gelder et al., 1991; Stevenson et al., 2014; Williams et al., 2004; Zhang et al., 2019). These studies have consistently shown that individuals with ASD experience significantly weaker McGurk effects than the neurotypical population. However, it is important to note that the McGurk effect focuses on the cognitive cost of processing conflicting auditory and visual information. In contrast, in normal speech contexts, listeners avoid integrating conflicting audiovisual speech information (Brang, 2019; Seijdel et al., 2023). Specifically, the incongruent pairing that is necessary to elicit McGurk fusion responses has been regarded as artificial and unnatural, showing limited features present in everyday speech (Van Engen et al., 2022) because face-to-face conversations yield congruent combinations of auditory and visual speech. Moreover, McGurk studies tend to examine audiovisual processing using isolated phonemes rather than complete words, casting further doubt on their applicability to natural speech. Consequently, tasks that use more naturalistic stimuli, like complete words and congruent audiovisual pairings, may be better able to clarify the role of visual information in everyday speech perception, thus advancing beyond the limited context of the McGurk effect.

While strong correlative research has identified the left pSTS as a region associated with the generation of the McGurk effect, only two studies to date have used causal methods (Beauchamp et al., 2010; Hickok et al., 2018). In a 2010 study, Beauchamp et al. reported that single pulse transcranial magnetic stimulation (TMS) applied to the left pSTS significantly reduced perception of the McGurk effect, providing strong evidence for the role of the pSTS in the generation of this illusion. TMS is a noninvasive brain stimulation method that involves the application of magnetic pulses to targeted brain areas that causes neurons to immediately depolarize thus injecting noise into ongoing processes (Hallett, 2000). This method has been effectively used to study the causal mechanisms underlying numerous cognitive and perceptual processes (Hallett, 2000; Rossini & Rossi, 2007; Walsh & Cowey, 2000). In Beauchamp et al.’s 2010 study, the authors conducted two separate experiments, each with sample sizes of 9, using similar task designs but two different speakers (experiment 1 used a female speaker and experiment 2 used a male speaker) and two different phonemes (experiment 1 used auditory /BA/ with visual /GA/ and experiment 2 used auditory PA with visual /NA/ or /KA/). The authors used two separate experiments with different phonemes and speakers to ensure the robustness of their findings across different speakers and stimuli combinations. Results showed that single-pulse stimulation to the left pSTS reduced the average frequency of fused percepts in McGurk conditions by 54% (experiment 1) and 21% (experiment 2) compared to stimulation applied to the control site. The authors also showed that only stimulation applied within 100 ms of the onset of the auditory stimulus (100 ms before until 100 ms after) reduced the frequency of McGurk percepts, while stimulation applied outside of this time range did not influence frequency.

Extending this research to clinical populations, Hickok et al. (2018) tested patients with a recent stroke using a McGurk effect paradigm with the goal of identifying lesioned areas of the brain that reduced McGurk effect percepts. Partially replicating the TMS work, Hickok et al. (2018) showed that stroke lesions in the broad superior temporal lobe (as well as in auditory and visual areas in the superior temporal and lateral occipital regions) resulted in the greatest deficits in McGurk perception, adding support to the model that the left pSTS enables the generation of the McGurk effect.

While both prior TMS and stroke lesion mapping studies identified the left pSTS as being causally relevant to the generation of the McGurk effect, those studies were not designed to test the relevance of this structure on more natural, congruent audiovisual speech perception behaviors. Building upon Beauchamp et al. (2010)’s findings, here we sought to address the involvement of left pSTS in other aspects of audiovisual speech processing beyond the generation of the McGurk percept and further investigate whether the McGurk effect is a good proxy for audiovisual processing. Based on past literature, two clear predictions emerged: 1) transient disruption of left pSTS activity will impair both the McGurk effect and the normal benefits from audiovisual speech. Such a finding would indicate that left pSTS is a critical hub for audiovisual speech generation in general and that the McGurk effect is a good proxy for natural audiovisual speech behaviors. Or 2) transient disruption of left pSTS activity impairs the McGurk effect with minimal impact on the normal benefits from audiovisual speech. Such findings would indicate that this region is likely only one of many critical structures necessary for audiovisual speech generation in general, reflecting only a subset of the information relayed from visual to auditory speech regions and laying the groundwork for research to understand which information is relayed through this hub.

To test these predictions, we assessed the impact of TMS on both the frequency of subjects’ McGurk effect percepts (which captures how visual information can change, or modulate, the perception of auditory information) and measures of audiovisual facilitation (which focuses on how visual information aids and facilitates the processing of the auditory information). By distinguishing between these two measures, we can better understand how mismatched visual information can modulate auditory perception and whether this is dissociable from the ability of congruent visual information to improve and facilitate the processing of concurrent auditory information. Towards this goal, we used an audiovisual task with real word stimuli, rather than phonemes that are typically utilized in most McGurk-type designs, as well as a larger sample (*n* = 21) than the prior TMS study. We hypothesized that pSTS stimulation would affect the McGurk effect more than congruent audiovisual benefits, providing evidence that the visual modulation of auditory speech relies on different neural mechanisms and consequently brain regions from audiovisual facilitation.

## Methods

### Subjects

25 healthy subjects (10 males, mean age = 24.7, 4 left-handed) with self-reported normal hearing and vision without a history of neurological disorder participated in the study. Four of the 25 total subjects either voluntarily withdrew from the study or experienced data errors resulting in 21 total subjects who completed the study (7 male, mean age = 24.2). Beauchamp et al. (2010) prescreened their subjects to include only those who reported strong McGurk effects. They justified this pre-selection process because there are large individual differences in susceptibility to the McGurk effect (Nath & Beauchamp, 2012) and variable reliance on lip movements during speech perception (Gurler et al., 2015). However, we did not exclude any subjects based on their McGurk susceptibility. While previous research has shown that the specific audiovisual stimulus used affects the strength and frequency of McGurk effects experienced by subjects (Basu Mallick et al., 2015), the stimuli used in the current study successfully evoked McGurk percepts in the majority of individuals tested in a prior study by our lab (Brang et al., 2020) and the same stimulus set has been shown to produce robust congruent audiovisual benefits during speech recognition (Ross et al., 2007). Therefore, to maximize the generalizability of our study to everyday speech perception, we did not exclude subjects based on their susceptibility to the McGurk effect.

To estimate the necessary sample size, we conducted an a priori power analysis using G*Power (Faul et al., 2007) based on Beauchamp et al. (2010)’s data. In comparing the frequency of reported fusion responses during pSTS stimulation versus control site stimulation, their results yielded Cohen’s D values of 3.22 and 8.43 across two experiments. Given the effect size of 3.22 (the smaller of the two estimates) is considered extremely large using Cohen’s criteria (1988), we would need a minimum sample size of 4 to replicate their effects with a significance criterion of alpha = .05 and power = .95. However, as our goal was to examine whether congruent audiovisual behaviors were affected as well, we made the more conservative assumption that if present, TMS effects on congruent audiovisual behaviors would be at least 25% the magnitude of effect of TMS on McGurk percepts. Repeating the power analysis with an alpha of 0.05, power of .80, and effect size of .805 in a two-tailed paired t-test design yielded a minimum sample size of 15. We sought to exceed this number and collect as much data as possible before the end of April 2023.

All participants gave informed consent prior to the experiment. This study was approved by the institutional review board at the University of Michigan.

### Experimental Task

Our audiovisual speech task used stimuli adapted from a prior study by Ross et al. (2007). From their larger set, we selected 32 single syllable words starting with the consonants ‘b’, ‘f’, ‘g’, or ‘d’, with the initial vowel sound approximately balanced across the consonant groups (e.g., ‘bag’, ‘gag’, ‘dad’, ‘fad’).

The schematic of the trials is shown in Figure 1. On each trial, a female speaker produced a high frequency monosyllabic word starting with the consonant ‘b’, ‘f’, ‘g’, or ‘d’. The trials were either audio-only, visual-only, audiovisual incongruent, or audiovisual congruent. Pink noise (SNR of -4.6 dB) was applied to all auditory stimuli (visual-only trials included pink noise but no speech) to increase the relative difficulty of the task, to avoid ceiling effects, and because the addition of noise tends to increase the reliance on visual speech information (Ross et al., 2007). The noise level was set based on piloting to lower accuracy in the audio-only condition away from the ceiling. In trials with a visual stimulus, the speaker’s video appeared 500 ms prior to the onset of the auditory stimulus. In audio-only trials, a gray box appeared 500 ms prior to the auditory onset to provide an equivalent temporal cue. Visual stimuli were recorded at 29.97 frames per second, trimmed to 1100 ms in length, and adjusted so that the first consonantal burst of sound occurred at 500 ms during each video.

**Figure 1.**
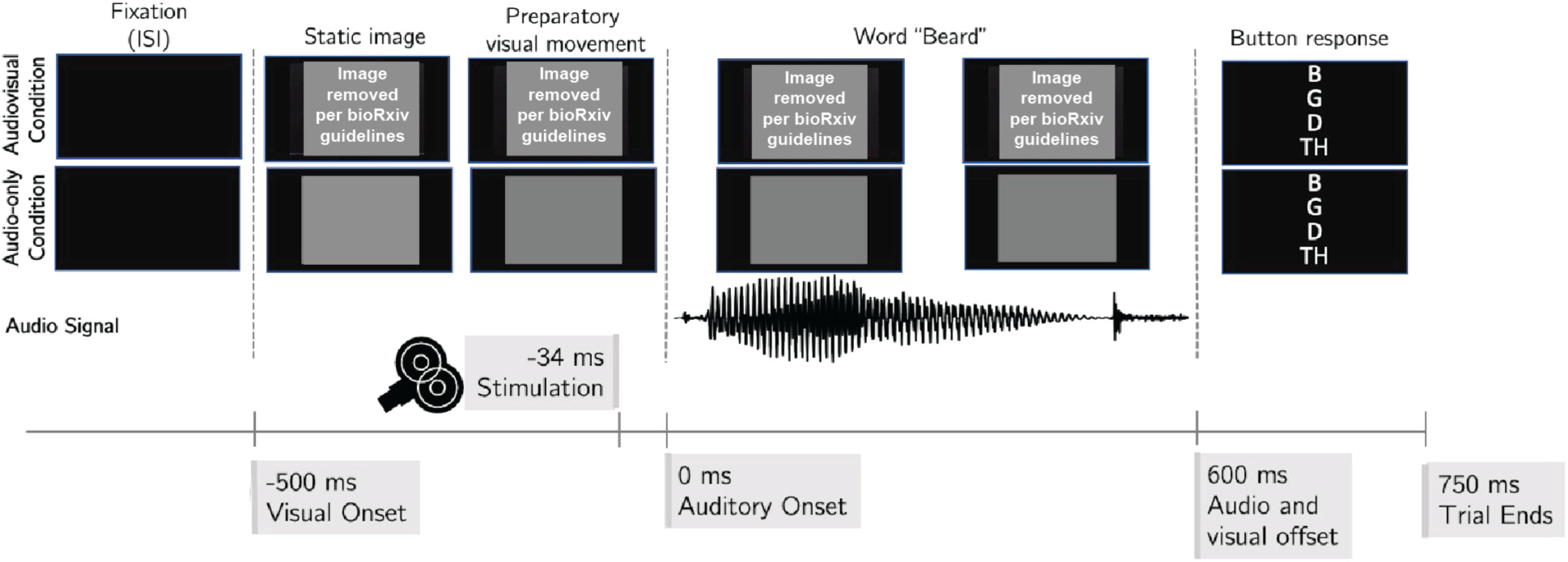
Trial schematic for the word ‘beard’. All trials started out with a black screen lasting between 125 - 375 ms. 500 ms prior to the auditory onset, either a gray box (audio-only condition) or the video of the speaker (AV and lipreading conditions) appeared. For blocks in which TMS was applied, stimulation was applied 34 ms prior to the onset of the audio. Following the audio/visual offset, participants were given 1.5 seconds to identify via button press which initial consonantal sound the word they heard. Note that each of the three conditions had pink noise mixed in with the audio signal (the visual-only condition contained only the pink noise at time 0).

In Beauchamp et al. (2010), the authors found maximally diminished fusion responses when the pSTS was stimulated 34 ms before auditory onset. Accordingly, here we applied a single TMS pulse 34 ms prior to audio onset (see TMS parameters below). Six hundred milliseconds following auditory onset, subjects were prompted to report the initial consonant of the seen (in visual trials) or heard word using a gamepad (Logitech F310) from the 4 options displayed on the screen. Subjects were asked to choose the option that sounded closest to what they heard if they were unsure. They were informed that the word that they heard may not be real words. The 4 response options always included the initial consonant of the spoken word, the initial consonant of the video (i.e., a viseme, which is the visual equivalent of a phoneme in spoken language), as well as the two common McGurk fusion percepts. These four response options remained consistent across all stimuli and conditions, even if some options were not relevant to certain conditions. The task was completed via a desktop computer using Psychtoolbox-3 (Brainard & Vision, 1997; Kleiner et al., 2007; Pelli & Vision, 1997) with participants seated approximately 60 cm away from the screen at eye level.

The task consisted of 3 blocks in total. Two blocks included TMS stimulation of one anatomical region per block (the left pSTS or vertex; defined below) and one block included no stimulation. The order of stimulation was counterbalanced across participants. In total, the study consisted of 384 trials, with 128 trials in each block with 4 audiovisual conditions (audio-only, visual-only, audiovisual congruent, audiovisual incongruent) divided equally within each block (32 trials per condition per block). Each block took approximately 7 minutes to complete, and participants were given the option to take breaks between each block.

Based on prior literature and internal piloting in which subjects provided free-response reports of what they heard, McGurk fusions were expected for two sets of audiovisual combinations: auditory words starting with either a B or F and visual words starting with either or G or D, respectively. For example, auditory ‘buy’ + visual ‘guy’ typically resulted in the percept ‘die’ or ‘thigh’, and auditory ‘fad’ + visual ‘dad’ typically resulted in the percept ‘tad’ or ‘thad’. To ensure that the auditory and visual components of each word were presented the same number of times throughout the experiment, half of the incongruent audiovisual trials had the modality of these pairings flipped (auditory words starting with either a G or D and visual words starting with either or B or F, respectively). These flipped pairings were not expected to generate fused responses, although they were still expected to reduce accuracy and slow reaction times; during piloting subjects were more likely to report ‘hearing’ the visual percept than a fused percept.

### MRI and TMS Procedure

Prior to the behavioral testing, on a previous day all participants underwent a structural MRI scan using a 3T GE MR 750 scanner to acquire T1-weighted images to be used for TMS guidance; three of 21 participants’ MRI data was acquired as part of a different experimental paradigm from our lab (Ganesan et al., 2022). Individual participants’ T1 scans were processed using Freesurfer (http://surfer.nmr.mgh.harvard.edu/) for cortical reconstruction and volumetric segmentation. The Freesurfer-generated pial and white matter reconstructions were then used to localize the left pSTS target coordinates according to the labels of the automatic cortical parcellation and automatic segmentation volumes. Specifically, we used the individual subject coordinates from the center of ‘lh-bankssts’, which is the bank of the left hemisphere superior temporal sulcus, as our pSTS target, and the vertex was defined as the midline of the postcentral gyrus using the subjects’ structural MRI scan.

TMS was applied through a MagPro X100 using a 70 mm figure-8 shaped TMS coil (MCF-B70, MagVenture Inc.). For each subject, their respective stimulation intensity was determined by obtaining their resting motor threshold and multiplying it by 1.1 (110%) as is the standard for single-pulse TMS thresholding (Kallioniemi & Julkunen, 2016; Sondergaard et al., 2021). The resting motor threshold is the lowest stimulation intensity necessary to evoke a consistent motor response while targeting the motor cortex. Specifically, this is the threshold at which an electromyographic motor response that is greater than 50 μV from the first dorsal interosseus muscle measuring is observed 5 out of 10 times (Mills & Nithi, 1997; Rossini & Rossi, 2007). In our motor thresholding, our study recorded from the right first dorsal interosseous muscle while stimulating the left primary motor cortex. The mean resting motor threshold used in our study was 51.7% of the maximum stimulator output. Once the target intensity was determined for each subject, the same intensity (110% of the resting motor threshold) was used for the entirety of their TMS session.

Following motor thresholding, participants completed the audiovisual task with each block targeting a different TMS site: pSTS, vertex, and no stimulation. The vertex stimulation block served as a control condition for the non-specific effects of TMS on behavior (e.g., scalp sensation, auditory stimulation, induced current in the brain, etc.) (Jung et al., 2016). Brainsight’s neuro-navigation system (Brainsight; Rogue Research) was used to target the stimulation sites in real time by registering participants’ facial landmarks to participants’ structural T1s using headbands containing trackers. The target coordinates for the pSTS that were obtained via Freesurfer reconstructions were used to guide TMS stimulation. Figure 2 shows the location of the left pSTS stimulation sites across subjects along with the anatomical label. The average coordinates of the left pSTS stimulation site across all subjects were (x = -65.1 ± 3.3, y = -48.5 ± 4.8, z = 7.3 ± 3.5) on a standard MNI-152 brain.

**Figure 2.**
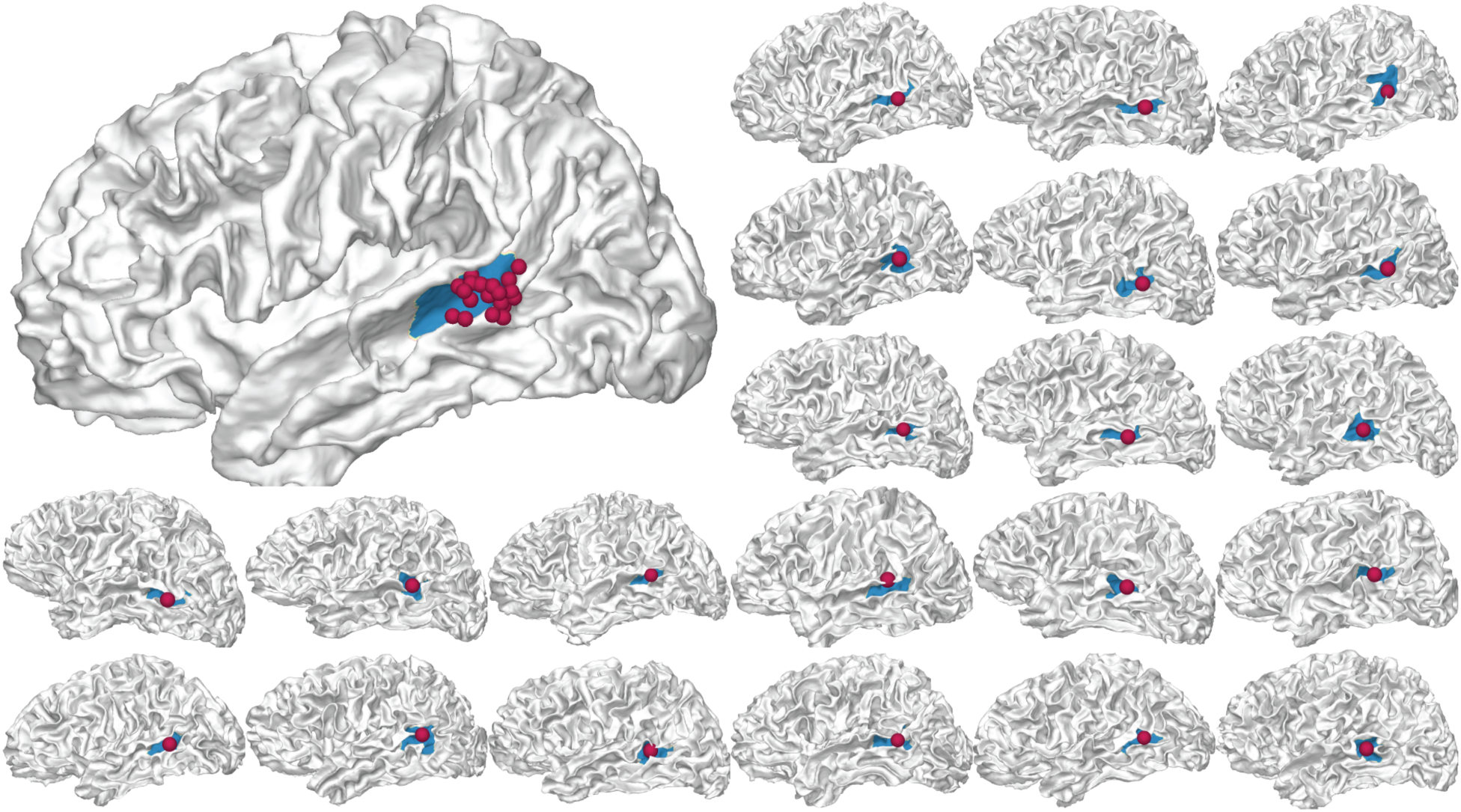
(top left) White matter rendering of the cvs_avg35_inMNI152 brain showing MNI transformed pSTS stimulation sites from all 21 subjects. Each sphere denotes the specific region where TMS was applied for a single subject. The blue region denotes the pSTS (Freesurfer label banksts) which was used for targeting. The smaller brains show the stimulation site for each of the 21 subjects on their own Freesurfer reconstructed brain.

To ensure precise delivery of the stimulation pulse relative to the onset of the auditory stimuli, the TMS pulse was auto-triggered through our MATLAB script using the MAGIC toolbox (Saatlou et al., 2018). Due to the position of the coil required to target the pSTS, earbuds, instead of headphones, were used to deliver the audio. Headband positions were validated both before and after the motor thresholding prior to the audiovisual task. If significant deviations from the original facial landmarks were observed, the landmarks were re-registered.

### Data Analyses

Our primary comparisons of interest examined changes in performance following stimulation of the left pSTS relative to Vertex stimulation for McGurk fusion frequency (the frequency at which subjects reported a fused percept on McGurk pairings) and for accuracy during congruent audiovisual trials. These measures were selected to be comparable with prior causal audiovisual research such as Beauchamp et al. (2010) and Hickok et al. (2018). As noted above, the incongruent audiovisual condition included 50% of stimuli combinations whose phonemic pairings typically evoke McGurk fusions (e.g., auditory ‘bet’ and visual ‘get’ yield the percept of ‘debt’) and the reversed pairings (e.g., auditory ‘get’ and visual ‘bet’) that do not typically result in McGurk fusions, for the purpose of counterbalancing stimuli. McGurk fusion frequency was only estimated from the 50% of incongruent audiovisual trials that contained auditory words starting with either a B or F and visual words starting with either or G or D, respectively. To ensure comparability with the McGurk fusion frequency measure, performance on congruent audiovisual trials was restricted to the same set of auditory words, unless noted otherwise. For example, the auditory word ‘bet’ was included in both the McGurk fusion frequency analysis (auditory ‘bet’ + visual ‘get’) and the congruent accuracy and RT analyses (auditory ‘bet’ + visual ‘bet’), but the auditory word ‘get’ was excluded (as auditory ‘get’ + visual ‘bet’ typically fails to yield fusion percepts). By restricting our analyses of congruent audiovisual trials, we ensured a similar base rate for the relevant comparisons across the conditions. Secondary analyses examined the overall pattern of results using a two-way ANOVA to test the impact of stimulation conditions (pSTS, vertex, or no stimulation) across the four conditions. These secondary analyses were conducted on all trials (McGurk and Non-McGurk stimuli combinations). Although the original degrees of freedom are reported here for clarity, p values were subjected to Greenhouse–Geisser correction where appropriate (Greenhouse & Geisser, 1959).

## Results

Figure 3 shows the accuracy and fusion response rates across the congruent, audio-only, and incongruent McGurk trials. As noted in the methods, McGurk fusion frequency was only estimated from the 50% of incongruent audiovisual trials that contained auditory words starting with either a B or F, and visual words starting with either or G or D, respectively. To ensure comparability with the McGurk fusion frequency measure, we first restricted our analyses to the same set of auditory words across all conditions (later analyses and Figure 4 reflect the data from all trials). Collapsing across stimulation site, congruent audiovisual trials showed higher accuracy relative to audio-only trials (t(20) = 14.4, *p* < .001, *d* = 3.14), and audio-only trials showed higher accuracy relative to incongruent McGurk trials (t(20) = 16.6, *p* < .001, *d* = 3.61) validating the positive and negative influence of vision depending on congruent and incongruent contexts.

**Figure 3.**
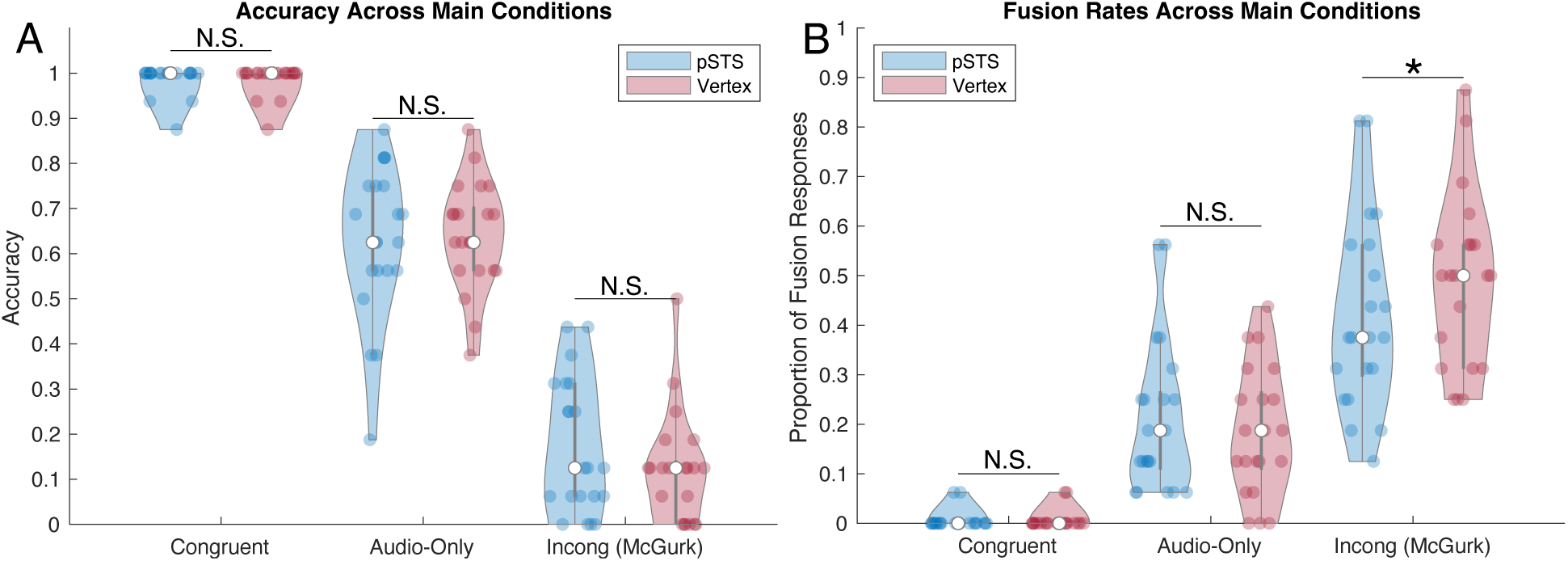
Violin plots showing accuracy (A) and fusion response rates (B) for pSTS and vertex stimulation sites, across the three main experimental conditions. Data were restricted to ‘McGurk’ stimuli to ensure comparability across analyses. Center circles indicate the median, gray boxes reflect the upper and lower quartile ranges, whiskers the min and max excluding outliers, and colored points are individual subject responses. TMS stimulation of the pSTS lowed the rate of fusion responses made by subjects on incongruent McGurk trials but did not impact congruent audiovisual trial accuracy. *p<.05.

**Figure 4.**
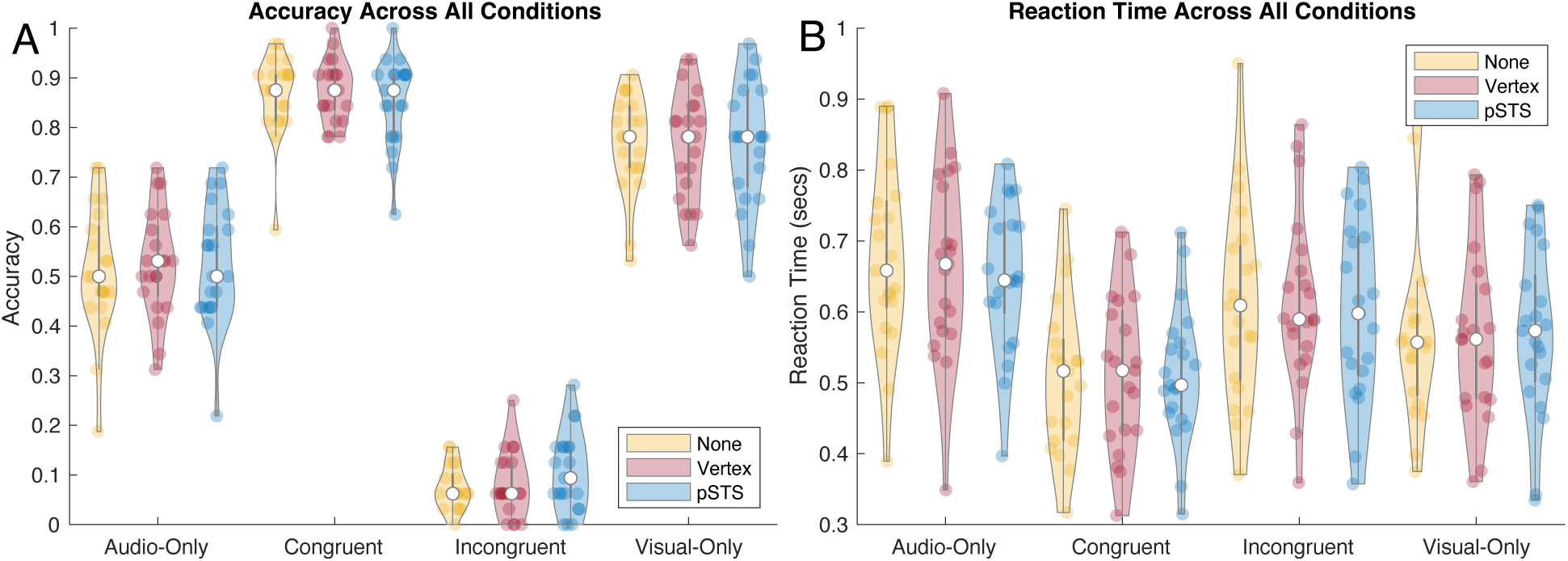
Violin plots showing accuracy (A) and reaction time (B) for each stimulation site and condition. Center circles indicate the median, gray boxes reflect the upper and lower quartile ranges, whiskers, the min and max excluding outliers, and colored points are individual subject responses.

To directly replicate the comparison made by Beauchamp et al. (2010), we examined the proportion of fusion responses made by subjects on Incongruent McGurk trials. Consistent with their data, single-pulse TMS to the left pSTS significantly reduced subjects’ likelihood of perceiving a McGurk fusion percept compared to the TMS to the vertex (*t*(20) = 2.16, *p* = 0.043, *d* = 0.472; Fig 3). Next, we examined the effect of TMS on congruent audiovisual accuracy. In contrast to McGurk trials, behavior on congruent audiovisual trials did not differ between stimulation of the pSTS and Vertex (t(20) = 0.00, *p* = 1.00, *d* = 0.00; Fig 3A); note that the statistics are at the absolute minimum because the average means across stimulation sites were identical. Comparing McGurk fusion rates and congruent audiovisual accuracy in a 2x2 repeated measure ANOVA additionally demonstrated a significant interaction between the two [*F*(1,20) = 5.15, *p* = .034, *ηp^2^* = 0.205] indicating that pSTS stimulation affected McGurk fusion to a greater degree than congruent audiovisual accuracy.

Following these planned comparisons, we calculated repeated measures ANOVAs for accuracy and reaction time (RT) data across all conditions. Violin plots showing the distribution of accuracy and reaction time measures are shown in Figure 4. Repeated measures ANOVA applied to accuracy data revealed a main effect of visual type [*F*(3,60) = 561.9, *p* = 7.4E-39, *η_p_^2^* = 0.913], but no effect of stimulation site [*F*(2,40) = 0.242, *p* = .737, *η_p_^2^* = 0.002], nor an interaction between visual type and stimulation site [*F*(6,120), *p* = .725, *η_p_^2^* = 0.007]. These results demonstrated a strong influence of visual content on task performance (such that congruent visual speech improves speech recognition and incongruent visual speech impairs it) and that there was no systematic effect of stimulation across conditions. Notably, visual only performance was higher than audio-only performance likely due to the high level of auditory-noise.

RT data mirrored those of the accuracy data with a main effect of visual type [*F*(2.3,46.08) = 54.379, *p* = 1.19E-13, *η_p_^2^* = 0.185], but no effect of stimulation site [*F*(2,40) = 0.152, *p* = 0.860, *η_p_^2^* = 0.001], nor an interaction between visual type and stimulation site [*F*(6,120) = 0.820, *p* = 0.556, *η_p_^2^* = 0.002]. Moreover, regardless of stimulation site, congruent visual speech was responded to more quickly than incongruent speech (*t*(20) = 8.44, *p* < .001, *d* = 1.84), which in turn was faster than audio-only trials (*t*(20) = 3.86, *p* < .001, *d* = 0.842) showing the strong benefit of visual information regardless of congruity. Stimulation of the pSTS (relative to vertex stimulation) affected neither the RTs of congruent trials (*t*(20) = 0.00, *p* = 1.00, *d* = 0.00) nor incongruent trials (*t*(20) = 0.552, *p* = .587, *d* = 0.121) showing the resistance of this audiovisual speeding to pSTS stimulation.

## Discussion

This study used real word stimuli to investigate the role of the left pSTS on audiovisual speech processes, including McGurk effect percepts, audiovisual facilitation, audiovisual reaction times, and lipreading. Using TMS in healthy subjects, our data demonstrate that stimulation of the left pSTS significantly disrupts the experience of the McGurk effect, reducing the frequency of reported fusion percepts, while leaving the other audiovisual processes intact. Our study extends the work of Beauchamp et al. (2010) demonstrating that TMS applied over the left pSTS, but not a control site (vertex), reduces the strength of the McGurk effect. However, as their behavioral task could not robustly capture the benefit of congruent visual stimuli on auditory speech perception, it remained unclear whether this region contributes specifically to McGurk processes, or to the general audiovisual speech integration processes at large. Our study extends Beauchamp et al. (2010)’s finding by confirming that the neural mechanism facilitating the McGurk perception, which many consider unnatural and artificial, can be dissociated from the other audiovisual processes like audiovisual facilitation and lipreading, which occur in everyday speech. Similarly, our results are consistent with models proposing that the pSTS is only one of the multiple critical areas supporting audiovisual speech interactions. This work adds to the growing evidence that McGurk processing relies on additional neural mechanisms beyond our everyday audiovisual speech.

One way to explain the involvement of pSTS in McGurk processing but not in audiovisual facilitation is that direct projections from visual motion area MT/V5 to the auditory cortex (Besle et al., 2008) allow the visual facilitation of auditory process to bypass the pSTS. Similarly, it is also possible that the neural processes underlying audiovisual facilitation alternatively recruit frontal structures to recover speech information through sensorimotor integration (Du et al., 2014, 2016; Hickok & Poeppel, 2007) without involving the pSTS. Along with Beauchamp et al. (2010)’s findings of reduced McGurk effect when targeting the left pSTS that we replicated here, a similar report of a weakened McGurk effect has been reported in patients following strokes near the left pSTS (Hickok et al., 2018). Adding onto these findings, our work provides compelling evidence that McGurk processing is a specific form of audiovisual speech integration that’s independent of other audiovisual speech enhancement. While McGurk processing may share some naturalistic features of everyday audiovisual speech processing, it also contains additional less ecologically valid properties like mismatched auditory and visual information. Consequently, it is possible that the left pSTS is more responsible for detecting or reconciling minor incongruities across modalities and re-evaluating the transformation of visemes to phonemes.

The work of Hickok et al. (2018) and Van Engen et al. (2017) similarly aimed to investigate the relationship between McGurk susceptibility and the use of visual information to facilitate speech perception. Indeed, both studies reported minimal correlations between the two measures, providing evidence against the widespread use of McGurk susceptibility as an index for audiovisual speech integration (Alsius et al., 2007; Jones & Callan, 2003; Paré et al., 2003; Van Wassenhove et al., 2007).

While we replicated Beauchamp et al. (2010)’s main finding, such that single pulse TMS to left pSTS diminishes the McGurk effect, we observed a much smaller change in behavior. In comparison to the large effect size reported by Beauchamp et al. (2010), with a Cohen’s D of 3.22 and 8.43 (across two separate experiments using different speakers and phonemes), our effect size for the difference in the frequency of fusion responses was much more moderate, yielding a Cohen’s D of 0.472. Such disparity in the effect size may be accounted for by the differences in the two study designs or the common trend for effect sizes to lower with larger sample sizes (e.g., Slavin and Smith (2009).

Our study differed from Beauchamp et al. (2010) in multiple ways. First, Beauchamp et al. (2010) only had 2 conditions: congruent and incongruent McGurk conditions. All trials used the same single auditory phoneme either matched with its congruent viseme or an incongruent viseme that is known to create a fusion percept. Conversely, we included multiple additional conditions to provide context for the subjects’ performance and to enable counter-balancing stimuli. Second, we used real monosyllabic words rather than phonemes for more naturalistic speech stimuli. This was done to address a common criticism of McGurk studies which argues that phonemes are highly artificial and do not reflect natural speech. Third, Beauchamp et al. (2010) determined the location of left STS using both anatomical (5 subjects based on landmarks) and functional (7 subjects based on individual subjects’ fMRI activation patterns) approaches. However, our study relied only on anatomical landmarks to determine the intended stimulation site. Fourth, we used the vertex as a control site rather than “a control TMS site dorsal and posterior to the STS” as reported by Beauchamp et al. (2010). This was to ensure we were using a consistent control site across subjects. Fifth, we did not exclude subjects based on whether they experienced strong McGurk effects. Our prior work (Brang et al., 2020) using similar stimuli set showed that most individuals report some level of fusion responses when presented with our word stimuli. Therefore, we wanted to ascertain that the results of our TMS were generalizable and not restricted to only those who experience strong McGurk percepts. Indeed, in the no-stimulation condition for the current dataset, all participants experienced a decrease in accuracy in the McGurk audiovisual incongruent condition relative to the audio-only condition (range 12.5 - 75% decrease in accuracy, mean = 45.5%). Sixth, we added pink noise to all our auditory stimuli whereas no noises were added to Beauchamp et al. (2010)’s auditory stimuli. Our decision to add noise was based on the prior literature showing that dependence on visual speech information increases with introduction of noise (Alsius et al., 2016; Buchan et al., 2008; Stacey et al., 2020). The addition of pink noise may in part have aided in generating McGurk percepts in our participants. Lastly, the two studies differed slightly in the TMS threshold used to apply single pulse stimulation. Whereas Beauchamp et al. (2010) used 100% of the resting motor threshold (RMT) for pulses, we used a slightly higher threshold at 110% of the RMT. We opted for the higher threshold as 110% or 120% of RMT is the more widely reported approach found in TMS literature (Cuypers et al., 2014; Kallioniemi & Julkunen, 2016; Sondergaard et al., 2021) which would naturally be expected to produce greater disruption of the involved region.

Given these differences in the study designs, it is possible that the disparity in the effect sizes of the pSTS stimulation on the frequency of McGurk effect may have been driven by a few of these factors. Specifically, we speculate that the largest driver of the disparity is the difference in the exclusion criteria (which were accompanied by other task designs to ensure that subjects still get fusion percepts). By broadening the subject pool, we similarly broaden our inference beyond the individuals who almost always experience the McGurk effect with a particular stimulus pairing. Because our pool of subjects is less likely to experience the McGurk percept compared to those from Beauchamp et al. (2010), it is possible that the effect of pSTS stimulation is less pronounced as the integration of the incongruent stimuli does not always occur as is the case for Beauchamp et al. (2010)’s subjects. In addition, it is also widely accepted that the likelihood of the McGurk effect varies largely across the stimuli used (Basu Mallick et al., 2015; Beauchamp et al., 2010). It is plausible that the audiovisual phoneme pairing used in Beauchamp et al. (2010) elicits a stronger McGurk percept compared to the audiovisual word pairings used in our study.

Taken together, our data points to evidence that audiovisual speech integration is not exclusively dependent on a single major hub in the left pSTS; instead, the left pSTS is more important for the generation of McGurk perception, resolving the conflict between auditory and visual information so that the information can be perceived as a single percept, rather than two mismatching percepts. While pSTS has been dubbed the multisensory hub of the brain and is indeed necessary for certain facets of multisensory perception, the importance of this region has likely been inflated due to the field’s heavy reliance on McGurk stimuli in the study of audiovisual integration. Indeed, our data provides converging and complementary evidence with the growing number of both behavioral and electrophysiological studies (Arnal et al., 2009; Arnal et al., 2011; Eskelund et al., 2011; Faivre et al., 2014; Fingelkurts et al., 2003; Lange et al., 2013; Palmer & Ramsey, 2012; Roa Romero et al., 2015) that point to the existence of multiple processing pathways and question the generalizability of McGurk perception to more naturalistic audiovisual speech processing. Additionally, the right pSTS or other contextual feedback mechanisms could have supported intact congruent audiovisual benefits after temporary disruption of the left pSTS. Collectively, this data is consistent with the emerging viewpoint that two distinct neural pathways underlie congruent audiovisual processes responsible for speech enhancements and incongruent audiovisual processing responsible for McGurk processing. Specifically, one that enhances the initial encoding of auditory information based on the information passed by the visual cues and another that modifies auditory representation based on the integrated audiovisual information. The first early feedforward process may occur early in the processing stream and align auditory encoding with the temporal and acoustic features of the accompanying visual input (Arnal et al., 2009; Arnal et al., 2011; Van Wassenhove et al., 2005). The later feedback process is engaged following the detection of mismatch between the auditory and visual cues, with the brain subsequently altering and adjusting the processing of the unisensory speech based on the combined audiovisual information (Arnal et al., 2011; Kayser & Logothetis, 2009; Olasagasti et al., 2015). Given that this late feedback process is facilitated by higher order areas like pSTS, the limited reliance of this pathway during congruent audiovisual processing can explain why stimulation to pSTS shows limited disruption on the audiovisual speech enhancement benefits. Importantly, however, future research should aim to identify a double dissociation of sites responsible for congruent facilitation versus McGurk effects.

Importantly, while this study revealed a significant interaction between stimulation site and congruity, such that pSTS stimulation affected the McGurk effect but not congruent audiovisual benefits, the overall pattern of results was weaker than expected and warrants future replications. In particular, accuracy in the congruent condition approached ceiling which makes it more difficult to detect TMS stimulation effects. Nevertheless, there was no effect of stimulation on the strong reaction time benefits present in the audiovisual conditions, emphasizing that the pSTS does not appear responsible for audiovisual speeding. Moreover, as this was a within-subject, within-session study in which we observed effects on McGurk fusion rates, we would expect to observe some changes in congruent audiovisual condition if they were present.

In summary, our data demonstrate that while TMS to the left pSTS can limit audiovisual speech integration and result in a weaker McGurk effect, it does not universally reduce the ecologically important benefits of congruent visual information on speech perception. This suggests a dissociation in neural mechanisms such that the pSTS reflects only one of multiple critical areas necessary for audiovisual speech interactions.

## Acknowledgements

This study was supported by NIH Grants R00DC013828 and R01DC020717. The authors report no conflicts of interest.

